# Third Trimester-Equivalent Alcohol Exposure Reduces Neurons in Males and Increases Vglut2⁺ Neurons with Reduced Intrinsic Bursting in Females in The Murine Dorsal Subiculum

**DOI:** 10.64898/2026.05.20.726671

**Authors:** Katalina M. Lopez, Hyemin Choi, Adeline Feng, Lourdes Cazares, Javier Kelly Roman, Glenna J. Chavez, Melissa Molina Garcia, Julian Jaramillo, C. Fernando Valenzuela

**Affiliations:** Department of Neurosciences, School of Medicine, University of New Mexico Health Sciences Center Albuquerque, New Mexico, U.S.A

**Keywords:** Ethanol, fetal, development, hippocampus, subiculum, glutamate, electrophysiology

## Abstract

Individuals with Fetal Alcohol Spectrum Disorders (FASDs) show reduced subicular volume, and preclinical studies compliment this by demonstrating that third-trimester-equivalent ethanol exposure induced apoptosis in corticolimbic regions, including the subiculum. The subiculum mediates hippocampal-cortical communication critical for long-term memory consolidation. Within the distal dorsal subiculum, a population of bursting neurons uniquely express VGLUT2 and they play a key role in memory processing. We hypothesized that third-trimester-equivalent ethanol exposure would reduce neuronal and VGLUT2+ cell density in the dorsal subiculum and reduce the excitability of bursting neurons, providing a mechanism for long-term memory impairments observed in FASD. To test this, postnatal day (P)7 mice received a subcutaneous injection of ethanol and long-term effects were assessed in adolescence (P35-62). Using transgenic mice with fluorescently labeled VGLUT2+ neurons, and immunohistochemistry we observed a significant reduction in neuronal density in males and an increase in VGLUT2+ cell density in females. Using whole-cell patch clamp electrophysiology, we observed a reduction in action potentials per burst in both sexes. Additionally, females showed reduced overall excitability, and a subset of neurons exhibited a shift to regular spiking. These findings suggest that development ethanol exposure disrupts subicular output by impairing burst firing, potentially weaking hippocampal-cortical communication and contributing to the cognitive deficits associated with FASD.

**Highlights:**

- Third-trimester ethanol targets VGLUT2+ neurons in the dorsal subiculum
- Ethanol reduced neuronal density in male dorsal subiculum
- Ethanol increases VGLUT2+ cell density in females
- Ethanol reduces action potential per burst in both sexes
- Females show reduced excitability and loss of bursting in some cells

## 1. Introduction

Fetal Alcohol Spectrum Disorders (FASDs) encompass a continuum of cognitive and behavioral impairments, including deficits in attention, memory, and executive functioning, as well as increased seizure susceptibility in some cases (Bell et al., 2010; Hoyme et al., 2016; Popova et al., 2023). The prevalence of FASDs is estimated at 1–5% among children in the United States (May et al., 2018). Although the hippocampus proper has been widely studied in this context, the subiculum (SUB), a key output structure of the hippocampal formation, has received comparatively limited attention. Importantly, SUB alterations are implicated in numerous neurological and psychiatric disorders associated with FASDs (Baset & Huang, 2024; Lopez et al., 2026).

Clinical studies report that adolescents or young adults with FASD or prenatal alcohol exposure exhibit reduced hippocampal volumes that include the SUB (Astley et al., 2009; Roediger et al., 2021; Willoughby et al., 2008). Complementary findings further suggest that smaller hippocampal volumes are associated with poorer performance on episodic memory tasks, as well as depressive symptoms (Gimbel et al., 2025; Nakhid et al., 2023; Solar et al., 2022). Preclinical studies have also demonstrated that the SUB is particularly vulnerable to third-trimester-equivalent acute alcohol exposure (TTAAE) in rodent and non-human primate models (reviewed in Lopez et al., 2026). Ethanol (EtOH) reduces neuronal activity, triggering the intrinsic apoptotic pathway across multiple brain regions, including the SUB and other components of the limbic memory system (Bird et al., 2020, 2021, 2023; Farber et al., 2010; Ikonomidou et al., 2000; Olney et al., 2002; Wozniak et al., 2004). This TTAAE-induced apoptotic neurodegeneration leads to persistent neuronal loss in adolescence and early adulthood within the dorsal SUB (dSUB) (Lopez et al., 2025; Smiley et al., 2023).

Studies have focused on the inhibitory subicular neurons, with one study demonstrating that EtOH exposure at postnatal day (P)7 in mice reduces parvalbumin+ interneurons in the dSUB, which persists into early adulthood (Sadrian et al., 2014). Building on these findings, our laboratory identified increased apoptosis in γ-aminobutyric acid (GABA)ergic interneurons in the dSUB following P7 EtOH exposure. However, despite this early cell loss, total GABAergic interneuron density was not altered in adolescence. We also found reduced action potential firing in fast-spiking interneurons following EtOH exposure, particularly in female mice (Lopez et al., 2025). Taken together, the results of these studies indicate that fast-spiking parvalbumin+ interneurons are important targets of TTAAE.

Despite increasing recognition of SUB vulnerability to developmental alcohol exposure, relatively little attention has been given to excitatory pyramidal neuron subtypes in this region. As a central component of the hippocampal network, the dSUB is organized along a proximal–distal axis, with gradients of cell types that are both genetically and functionally distinct (Cembrowski et al., 2018). The dSUB receives major excitatory input from CA1 pyramidal neurons and projects broadly to cortical and subcortical targets, playing a key role in memory consolidation. Notably, vesicular glutamate transporter (VGLUT) isoforms are differentially expressed along the proximal–distal axis. VGLUT1 is enriched in proximal regions and is associated with regular-spiking pyramidal neurons, whereas VGLUT2 is enriched in distal regions and is associated with bursting pyramidal neurons (Wozny et al., 2018). These bursting neurons typically generate 2–3 action potentials per burst and are preferentially engaged during sharp wave ripple events, which are critical for memory replay and consolidation (Nitzan et al., 2020). During these events, bursting neurons exhibit reduced inhibition from local interneuron networks (Böhm et al., 2015).

Importantly, VGLUT2+ neurons in the distal dSUB project directly to layers II/III of the retrosplenial cortex. The CA1→dSUB→retrosplenial cortex circuit has been identified as a key pathway for initiating ripple propagation, underscoring the importance of distal dSUB VGLUT2+ neurons in memory-related network dynamics (Nitzan et al., 2020). Given the vulnerability of the dSUB to TTAAE and the specialized role of VGLUT2+ neurons in learning and memory, we hypothesized that such exposure would reduce the density of VGLUT2+ pyramidal neurons during adolescence and impair their intrinsic firing properties.We combined immunohistochemistry with *ex vivo* whole-cell patch-clamp electrophysiology to test this hypothesis.

## 2. Methods and Materials

### 2.1 EtOH exposure paradigm

All animal procedures were approved by the UNM-HSC Animal Care and Use Committee (protocol # 24-201560-HSC) and followed NIH guidelines. Transgenic VGlut2-ires-cre female mice (Jax 016963) crossed with B6.Cg-Gt (ROSA) 26Sortm3 (CAG-EYFP)Hze/J male mice (Jax 007903) were used to visualize VGLUT2+ cells in all studies. Mice were group-housed with same-sex littermates on a 12-h reverse light/dark cycle (lights on at 8 p.m.) and had *ad libitum* access to water and chow (Teklad 29290x diet, Envigo, Indianapolis, IN). P7 pups were randomly assigned to either the EtOH or saline treatment groups. The EtOH group received two subcutaneous injections of a 20% (v/v) EtOH solution in saline at a dose of 2.5 mg/kg, administered two hours apart, while control animals received saline alone. This injection regimen consistently produces peak blood EtOH concentrations of 400–500 mg/dL (Wozniak et al., 2004).

### 2.2 Immunohistochemistry and Unbiased Stereology

For quantification of VGLUT2+ cell density and neuronal density, mice received subcutaneous injections of ketamine (250mg/kg) and xylazine (25 mg/kg), were perfused transcardially with paraformaldehyde, and their brains were extracted as described previously (Bird et al., 2023). Sections (50 µm) from adolescent mice (P35-58) (corresponding to Bregma −2.69 to −3.79 in the adult mouse (Paxinos & Franklin, 2019)) were stained using both a rabbit IgG polyclonal anti-GFP (1:2000 dilution, reference #A11122, Life Technologies, Eugene, OR) and mouse IgG1 monoclonal anti-NeuN antibody (1:500 dilution, MAB377, Millipore Sigma, Burlington, MA). Prior to 24-hour incubation of those antibodies in 4°C, sections were incubated for 2 hours in blocking/permeabilization solution containing 1% bovine serum albumin, 5% donkey serum (Jackson ImmunoResearch, West Grove, PA) and 0.2% Triton X-100. After 24-hours, sections were incubated, at room temperature, in secondary donkey anti-rabbit IgG Alexa Flour 555 antibody (1:1000 dilution, catalog #A-31572, ThermoFisher) and a donkey anti-mouse IgG Alexa Flour 647 antibody (1:1000 dilution, #A-31571, ThermoFisher) for 2 hours. Sections were stained with 4′,6-diamidino-2-phenylindole (DAPI) and mounted as previously mentioned (Bird et al., 2023). Sections were imaged using a Zeiss Axioscan Z1 (Preclinical Core, Center for Brain Recovery and Repair, UNM-HSC) at 20X magnification. Images for the dSUB from both hemispheres were collected ∼15-20 sections per animal), and fluorescence channels included blue (DAPI for nuclei), orange (anti-GFP tagged VGLUT2+ pyramidal neurons) and red (anti-NeuN tagged neurons). Image processing and analysis were conducted using QuPath (Bankhead et al., 2017) by a blinded investigator. The dSUB was manually outlined, and we then used a script to divide it into 8 distinct regions along the proximal-distal axis. The densities of VGLUT2+ and NeuN+ cell densities were then automatically detected using a threshold of fluorescence detection by area (per mm^2^). A second blinded investigator counted the same brain section, and the average of both evaluators’ counts was used to ensure consistency and reduce human error.

### 2.3 Whole-Cell Patch Clamp Electrophysiology

Mice (P39-62) were anesthetized with a subcutaneous injection of ketamine (250 mg/kg) and xylazine (25 mg/kg). The brains were removed and placed in an ice-cold protective cutting solution for 2 minutes (containing in mM: 163.9 sucrose, 5 ascorbic acid, 10 MgSO_4_, 0.5 CaCl_2_, 2.5 KCl, 1.25 NaH_2_PO_4_, 30 NaHCO_3_, 20 HEPES, 25 glucose, 2 thiourea, and 3 sodium pyruvate, bubbled with 95%O_2_/5%CO_2_; pH 7.3-7.4, 300-310 mOsm, adjusted with KOH and HCl). Coronal sections (250 μm) containing the dSUB were cut using a vibratome (Leica VT10000 S, Leica Microsystems). Slices were transferred to a temperature-controlled holding chamber (Model 7470, Campden Instruments, Loughborough, England) containing a recovery solution (concentration in mM: 93.7 NaCl, 2.5 KCl, 1.25 NaH_2_PO_4_, 30 NaHCO_3_, 20 HEPES, 25 glucose, 2 thiourea, 3 sodium pyruvate, 5 ascorbic acid, 1 MgSO4, and 2 CaCl2, saturated with 95% O2/5% CO2, pH 7.3-7.4, 300-310 mOsm). The temperature was gradually reduced from 34°C to ∼21°C over a 90-minute recovery period. Recordings were performed in a chamber with a slice support (Model RC-27L, Warner Instruments), continuously perfused with artificial cerebral spinal fluid (ACSF) (containing in mM: 125 NaCl, 2 KCl, 1.3 NaH2PO4, 26 NaHCO3, 10 glucose, 1 MgSO4, 2 CaCl2, and 0.4 ascorbic acid; saturated with 95% O2/5% CO2, pH 7.3-7.4, 300-310 mOsm), delivered at 2 mL/min via a peristaltic pump (Master Flex Model 7518-10, Cole-Palmer, Vernon Hills, IL). Slices were held in place with a platinum wire (Catalog #43288, Alfa Aesar) and viewed using an Olympus BX51W1 microscope with a U-PMYVC camera adapter and lenses (Plan 4X, 0.10 NA; LUMPlan F1/IR 40X water immersion, 0.8 N.A., Olympus). EYFP+ cells in the dSUB were identified using a YFP filter cube (excitation 513-514 nm; emission 527-529 nm) illuminated by a mercury arc lamp. Recording electrodes (2-6 MΩ) were pulled from filamented borosilicate glass capillaries (Catalog #BF150-86-10) using a DMZ Universal Puller (Zeitz-Instruments Vertriebs GmbH, Martinsried, Germany) and controlled using an MP-225 micromanipulator (Sutter Instrument, Novato, CA) they were then filled with a K-gluconate based internal solution (concentration in mM: 135 K-gluconate, 6 KCl, 10 HEPEs, 2 MgCl_2_, 0.2 EGTA, 2 Na_2_-ATP, 0.5 Na_2_-GTP, 5 Na_2_-phosphocreatine, 0.2% Biocytin, pH 7.3-7.4, 290 mOsm) that created a liquid junction potential of −23.1 mV with the ACSF. The liquid junction potential was corrected by holding the membrane potential at −93.1 mV during whole-cell current-clamp recordings, yielding an effective holding potential of −70 mV. The recordings consisted of 15 steps of 500 ms current injections, starting from −200 pA in 100-pA increments. Recordings were amplified using an Axopatch 200B amplifier and digitized via a Digidata 1550B system, with Clampex software (v11, Molecular Devices). Traces showing a variation in access resistance>20% were excluded. Final analysis was performed offline using Clampfit v11 (Molecular Devices).

### 2.4 Statistical Analysis

Statistical analyses were performed using Prism version 10.6.1 (Graph-Pad Software, San Diego, CA) and R Statistical Software v4.4.0 (R Core Team, 2024). All data are presented as mean ± SEM. For each experiment, the unit of determination (*n*) was a single animal, sampled across 3-5 litters. Action potential recordings that were collected from multiple neurons within a single animal were averaged. Outliers were removed using a ROUT 1% test in Prism. Normality was also assessed in Prism using the Shapiro-Wilk test; data that did not pass were log-transformed and reanalyzed. Levene’s test was used to assess homogeneity of variance with the car package (Fox & Weisberg, 2019) in R. If the data passed the Shapiro-Wilks and Levene’s tests, the raw or log-transformed data were then analyzed using three-way ANOVAs with sex, region, and EtOH exposure as between-subject factors in Prism. Two-Way ANOVAs were then conducted with region and EtOH exposure as between-subject factors, separately for males and females. When ANOVA assumptions were not met by either normality or homogeneity of variance, data were analyzed using nonparametric aligned rank transform (ART) ANOVA using the ARTool v0.11.1 (Kay et al., 2025) and the readxl (Wickham & Bryan, 2025) packages in R. For electrophysiological measures collected across multiple current injection steps, repeated measures (RM) ART ANOVA models were used. For single-value outcomes (membrane resistance and rheobase), two-way ART ANOVA models without RM models were used. Planned comparisons were conducted to evaluate the specific effects of ethanol exposure for each sex on each outcome measure. Pairwise comparisons between control and ethanol-treated groups were performed using the ART-C contrast testing procedure, with Šidák correction (Kay et al., 2025).

## 3. Results

### 3.1 EtOH Exposure at P7 decreases total neuronal density in the dSUB of males but increases VGLUT2+ cell density in females at P35-58

We investigated whether EtOH exposure P7 produces long-term changes in neuronal density within the dSUB. Because VGLUT2 expression forms a gradient along the proximo–distal axis of the dSUB, the region was divided into eight segments along this axis to capture the gradient of expression (Cembrowski et al., 2018) (**Figure 1**).

**Figure 1.**
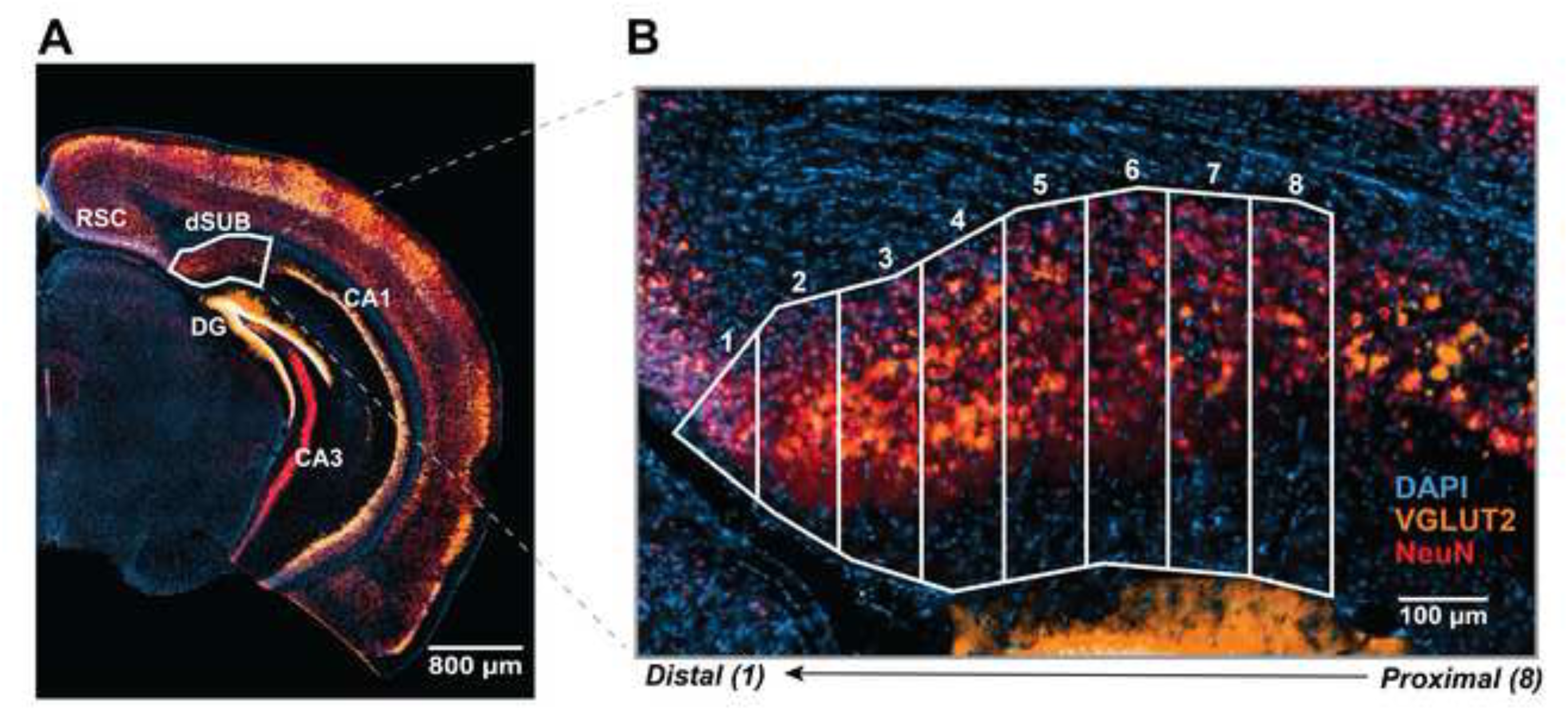
Regions of interest within the dorsal subiculum. **A)** Coronal section illustrating the dorsal subiculum, outlined in white. Scale bar = 800 μm. This image roughly correlates to Bregma −3.15 mm (Paxinos & Franklin, 2019). **B)** Higher-magnification of sample image of the dSUB showing the eight ROIs analyzed in this section using QuPath. Sections were stained with DAPI (blue), NeuN (red), and VGLUT2 (orange). Regions were defined along the proximal-distal axis, with region 1 representing the most distal and region 8 the most proximal location relative to CA3. Scale bar = 100 μm. Acronyms: dSUB = dorsal subiculum; RSC = retrosplenial cortex; DG = dentate gyrus; CA = cornu ammonis.

Quantification of neuronal density in the dSUB is shown in **Figure 2**. EtOH exposure reduced NeuN+ neuronal density in adolescent males but not females (**Figure 2B–C**). A three-way ANOVA revealed significant effects of region (*F*(7,279)=8.5), *p*<0.0001) and sex (*F*(1,279)=11.9), *p*=0.0006). Therefore, neuronal density was analyzed separately by sex. In females, total NeuN+ neuronal density was not altered by EtOH exposure (Two-way ANOVA: Treatment (*F*(1,136)=1.9, *p*=0.17). In contrast, males exposed to EtOH exhibited a significant reduction in NeuN+ neuronal density within the dSUB (Two-way ANOVA: Treatment (*F*(1,144)=11.4), *p*=0.0005). Both males and females showed significant regional gradients in neuronal density, with higher densities observed in distal regions (Two-way ANOVA (Region); Females (*F*(7,136)=3.7, *p*=0.001, Males (*F*(7,144)=4.7, *p*<0.0001) (**Supplementary Table 1**). To quantify VGLUT2+ neurons, we used a transgenic mouse line that expresses YFP in these neurons and detected it with an anti-GFP antibody to amplify the signal (**Figure 3A**). As in the NeuN analysis, the dSUB was divided into eight regions along the proximo–distal axis. A three-way ANOVA with Geisser-Greenhouse correction for unequal variances (**Supplementary Table 1**) revealed significant effects of sex (*F*(1,35)=4.4), *p*=0.043), region (*F*(1.9,61.5)=29.7, *p*<0.0001), and a treatment × sex interaction (*F*(1,35)=5.4, *p*=0.03) (Geisser-Greenhouse’s epsilon=0.27). Therefore, VGLUT2+ cell density was analyzed separately by sex. A significant effect of treatment was only observed in females (*F*(1,17)=7.9), *p*=0.01) (Geisser-Greenhouse’s epsilon=0.28) (**Figure 3B**). In contrast, significant region effects were observed in both females (*F*(1.9,30.2)=12.01, *p*=0.0001)(Geisser-Greenhouse’s epsilon=0.28) and males (*F*(7,136)=9.5, *p*<0.0001) (**Figure 3C**). This cohort included 39 mice from 16 litters (Female control, n = 10 mice from 4 litters; female EtOH, n = 9 mice from 5 litters; male control, n = 10 mice from 5 litters; male EtOH, n = 10 mice from 5 litters). Overall, these results indicate that P7 EtOH exposure increases VGLUT2+ cell density along the proximo-distal axis of the dSUB of females as well as reduces total neuronal density in males, consistent with our previous work (Lopez et al., 2025).

**Figure 2.**
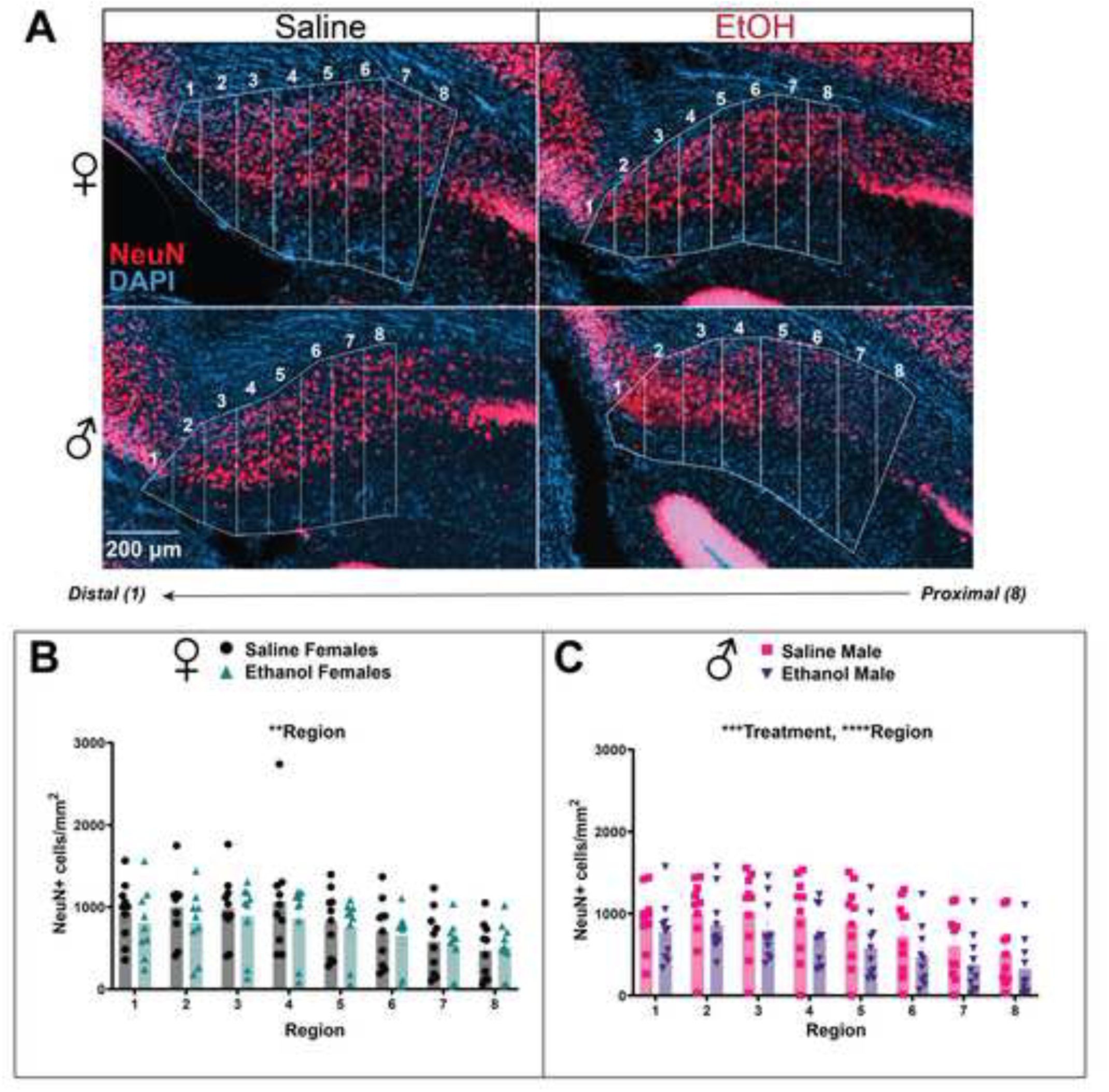
P7 ethanol exposure reduces total neuronal density in the dorsal subiculum of males during adolescence. A) Representative images of NeuN immunostaining (red) and DAPI (blue) from female (top) and male (bottom) saline control (left) and EtOH-treated (right) mice. Numbers (1–8) indicate positions along the proximodistal axis of the dorsal subiculum, with region 1 representing the most distal and region 8 the most proximal location. Scale bar = 200 μm. B–C) Summary graphs illustrating the effect of P7 ethanol exposure on NeuN+ neuronal density across dorsal subicular regions in females (B) and males (C). Data were first analyzed using a three-way ANOVA including sex, and region, which revealed significant effects of sex (*p* = 0.0006), region (*p* < 0.0001). Subsequent two-way ANOVAs performed separately by sex revealed a significant effect of region in females (\*\**p* = 0.001). In males, two-way ANOVA revealed significant effects of treatment (\*\*\**p* = 0.0005) and region (\*\*\*\**p* < 0.0001). Female control, *n* = 10 mice from 4 litters; female EtOH, *n* = 9 mice from 5 litters; male control, *n* = 10 mice from 5 litters; male EtOH, *n* = 10 mice from 5 litters.

**Figure 3.**
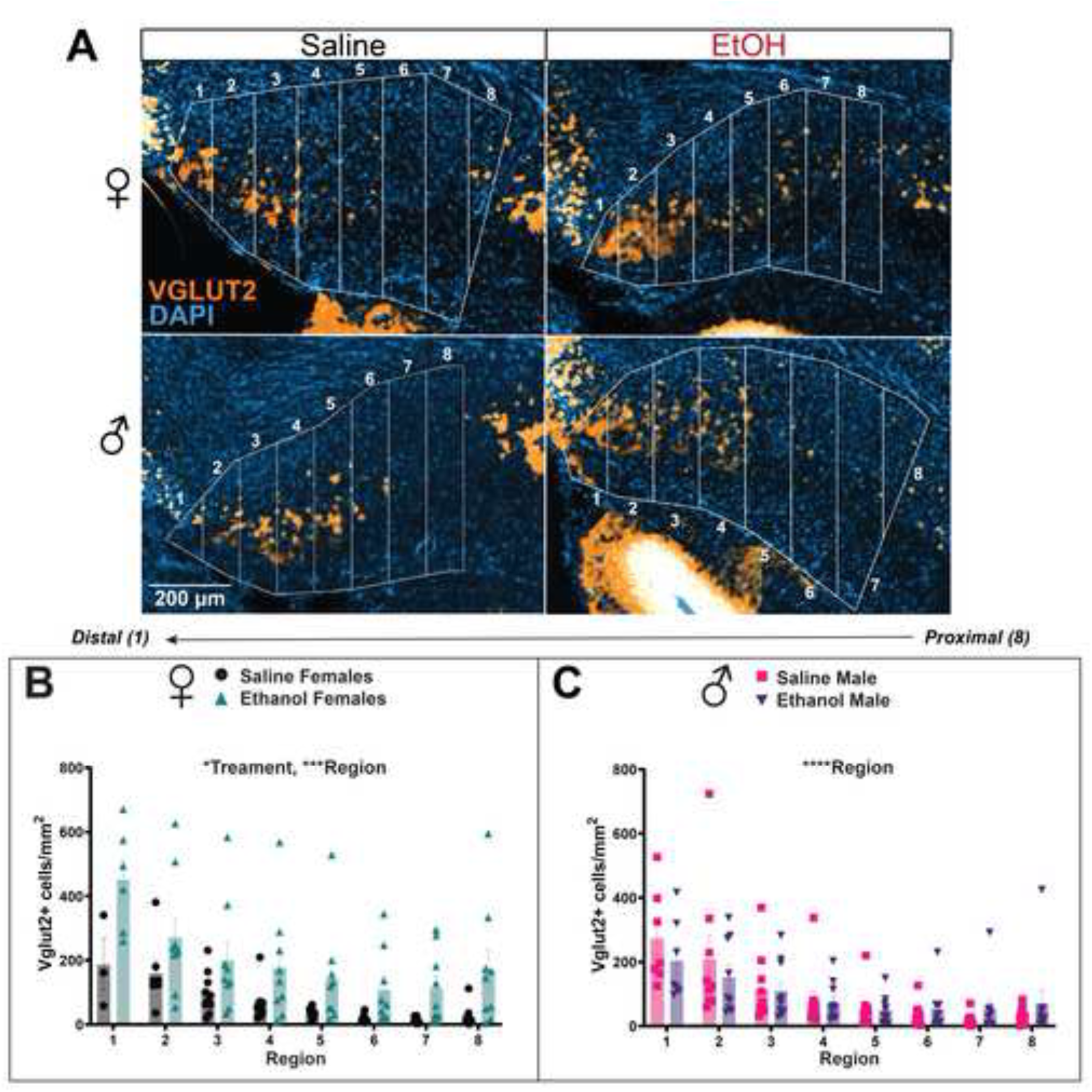
P7 ethanol exposure increases VGLUT2+ cell density in the dorsal subiculum in adolescent females. A) Representative images of VGLUT2 immunostaining (orange) and DAPI (blue) from female (top) and male (bottom) saline control (left) and EtOH-treated (right) mice. Scale bar = 200 μm. B–C) Summary graphs showing VGLUT2+ cell density across dorsal subicular regions in females (B) and males (C). Data were first analyzed using a three-way ANOVA including sex, treatment, and region, which revealed significant effects of sex (*p* = 0.042), region (*p* < 0.0001), and a treatment × sex interaction (*p* = 0.03). Subsequent two-way ANOVAs performed separately by sex revealed a significant effect of treatment (\**p* = 0.01) region in females (\*\*\**p* = 0.0001). Region was also a significant factor in males (\*\*\*\**p* < 0.0001), whereas treatment had no significant effect. Female control, *n* = 10 mice from 4 litters; female EtOH, *n* = 9 mice from 5 litters; male control, *n* = 10 mice from 5 litters; male EtOH, *n* = 10 mice from 5 litters.

### 3.2 EtOH exposure at P7 decreases action potential firing in burst-spiking subicular VGLUT2+ pyramidal neurons in adolescent females

We next investigated whether P7 EtOH exposure produces long-lasting alterations in the firing properties of VGLUT2+ bursting neurons (**Figure 4**). Using the YFP fluorescence, VGLUT2+ neurons were visually identified and randomly selected for whole-cell patch-clamp recordings. All recorded neurons were located in the distal dSUB, where VGLUT2 expression predominates. This electrophysiological cohort included 36 mice from 10 litters (Female control, n = 9 mice from 4 litters; female EtOH, n = 9 mice from 5 litters; male control, n = 9 mice from 3 litters; male EtOH, n = 9 mice from 4 litters). Passive membrane properties—including membrane capacitance, membrane resistance, and access resistance—were first assessed and did not differ by treatment or sex (**Table 1; Supplementary Table 1**).

**Figure 4.**
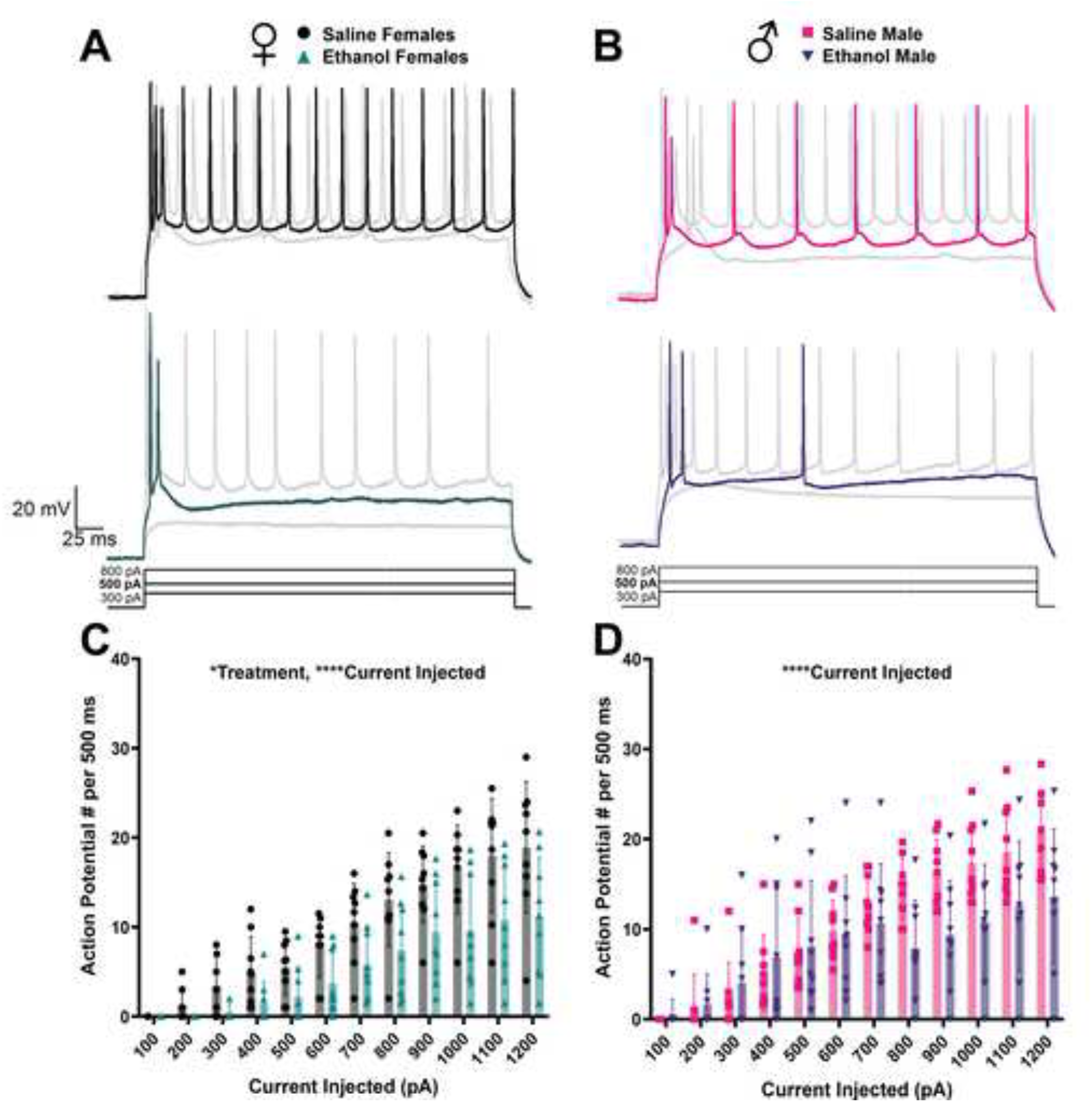
P7 ethanol exposure reduces action potential firing in VGLUT2+ bursting neurons in females. A) Representative traces of action potentials evoked by current injection (300, 500, and 800 pA) from saline (top, black) and EtOH-treated female mice (bottom, turquoise). B) Representative traces from saline (top, pink) and EtOH-treated male mice (bottom, purple). The central traces correspond to the 500-pA current step. Scale bar = 25 ms × 20 mV. C) Summary graph showing the number of action potentials generated per 500 ms current injection steps in female VGLUT2+ neurons from saline (black circles) and EtOH-treated mice (turquoise triangles). Two-way ANOVA revealed significant effects of treatment (\**p* = 0.01) and current injected (\*\*\*\**p* < 0.0001). D) Equivalent analysis in males showing saline (pink squares) and EtOH-treated (purple triangles) groups. Two-way ANOVA revealed a significant effect of current injected (\*\*\*\**p* < 0.0001) but no effect of treatment. Female control, *n* = 9 mice from 4 litters; female EtOH, *n* = 9 mice from 5 litters; male control, *n* = 9 mice from 3 litters; male EtOH, *n* = 9 mice from 4 litters.

**Table 1.** Effect of P7 EtOH exposure on passive membrane properties of dorsal subicular VGLUT2+ pyramidal neurons at P41-62. Membrane capacitance and access resistance data were analyzed by Two-Way ANOVA while Rm was analyzed by ART Two-Way ANOVA. There were no significant effects in treatment or sex. Values in parenthesis indicate the number of animals, from the total number of litters.

|  | Females |  | Males |  | P Values |
| --- | --- | --- | --- | --- | --- |
|  | Saline (9,4) | EtOH (9,5) | Saline (9,3) | EtOH (9,4) |  |
| Membrane Capacitance (pF) | 139.8±<br>49.32 | 147.6±<br>47.86 | 146.9±<br>25.13 | 156.2±<br>32.56 | Treatment P=0.56<br>Sex P=0.35<br>Interaction P=0.99 |
| Membrane Resistance (MΩ) | 115.9±<br>56.07 | 71.06±<br>28.54 | 108.4±<br>42.26 | 126.9±<br>57.85 | Treatment P=<br>0.37<br>Sex P= 0.19<br>Interaction P= 0.08 |
| Access Resistance (MΩ) | 19.33±<br>3.881 | 18.06±<br>4.007 | 17.64±<br>4.032 | 19.07±<br>3.484 | Treatment P=0.95<br>Sex P=0.80<br>Interaction P=0.30 |

Action potential firing was then analyzed across 500 ms current injection steps (**Figure 4A–B**). A three-way RM ANOVA revealed significant effects of injected current (*F*(1.7,55.5)=75.1, *p*<0.0001), sex (*F*(1,32)=11.04, *p*=0.002), and a current × sex interaction (*F*(1.7,55.5)=5.3, *p*=0.01). Therefore, analyses were conducted separately by sex. Adolescent females exposed to P7 EtOH exhibited an overall reduction in action potential firing (Two-way RM ANOVA; *F*(1, 16)=7.5, *p*=0.01). In contrast, males did not show a treatment effect (Two-way RM ANOVA; *F*(1,16)=3.6, *p*=0.07).

Additional action potential properties were also examined, including instantaneous firing frequency, overshoot amplitude, peak amplitude, threshold, and rheobase (**Table 2**). For instantaneous frequency, a three-way ART RM ANOVA revealed significant effects of sex (*F*(1, 275)=10.05, *p*=0.002), current injected (*F*(9,275)=3.8, *p*=0.0001), and a treatment × sex interaction (*F*(1,275)=6.9, *p*=0.009). However, when analyzed separately by sex, no significant treatment effects were observed; only the effect of current injected remained significant in females (Two-way RM ART ANOVA; *F*(11, 144)=2.5, *p*=0.007) and males (Two-way RM ART ANOVA; *F*(9,133)=2.4, *p*=0.01).

**Table 2.** Effect of P7 EtOH exposure on action potential parameters of dorsal subicular VGLUT2+ pyramidal neurons at P41-62. The instantaneous frequency, overshoot and peak amplitude, as well as the threshold of the first action potential on the 700 pA current step and rheobase were analyzed by ART ANOVAs. Values in parenthesis indicate the number of animals, from the total number of litters.

|  |  | Females |  | Males |  | Significant<br>P values |
| --- | --- | --- | --- | --- | --- | --- |
|  | Injected<br>Current (pA) | Saline (9,4) | EtOH (9,5) | Saline (9,3) | EtOH (9,4) |  |
| Instantaneous<br>Frequency (Hz) | 100 | - | - | - | 12.35 ± 0 | Sex P=0.002<br>Current Injected<br>P=0.0001<br>Treatment x Sex<br>P= 0.009 |
|  | 200 | 71.05 ± 19.13 | - | 39.89 ± 16.86 | 11.94 ± 3.57 |  |
|  | 300 | 43.80 ± 11.04 | 92.14 ± 8.36 | 31.40 ± 0 | 40.41 ± 8.38 |  |
|  | 400 | 57.45 ± 8.58 | 85.15 ± 15.58 | 27.57 ± 17.94 | 49.78 ± 13.45 |  |
|  | 500 | 60.71 ± 9.51 | 69.57 ± 7.85 | 53.08 ± 6.52 | 43.70 ± 7.24 |  |
|  | 600 | 48.90 ± 9.91 | 43.28 ± 9.93 | 33.24 ± 2.06 | 41.56 ± 6.85 |  |
|  | 700 | 39.27 ± 4.43 | 59.93 ± 11.06 | 32.77 ± 2.00 | 38.70 ± 5.23 |  |
|  | 800 | 41.88 ± 1.42 | 53.15 ± 9.51 | 33.60 ± 2.47 | 38.41 ± 8.40 |  |
|  | 900 | 43.03 ± 2.50 | 38.70 ± 3.69 | 36.26 ± 2.97 | 41.44 ± 8.26 |  |
|  | 1000 | 44.78 ± 1.73 | 48.02 ± 8.55 | 37.84 ± 3.36 | 45.90 ± 8.35 |  |
|  | 1100 | 50.92 ± 2.39 | 53.88 ± 9.17 | 39.59 ± 3.81 | 48.86 ± 7.62 |  |
|  | 1200 | 54.22 ± 3.17 | 65.17 ± 8.38 | 41.83 ± 2.10 | 47.95 ± 5.54 |  |
| Overshoot<br>Amplitude (mV) | 100 | - | - | 16.72 ± 0 | -7.16 ± 0 | Current Injected<br>P <0.0001<br>Treatment x<br>Current Injected<br>P=0.001<br>(* ART-C<br>contrast testing<br>with Sidak<br>correction<br>p=0.049) |
|  | 200 | 31.29 ± 4.75 | - | 30.43 ± 10.40 | 20.08 ± 13.94 |  |
|  | 300 | 35.13 ± 2.91 | 33.87 ± 1.73 | 39.37 ± 6.01 | 25.92 ± 8.41 |  |
|  | 400 | 39.40 ± 4.29 | 29.02 ± 4.61 | 34.65 ± 4.99 | 34.10 ± 7.05 |  |
|  | 500 | 40.49 ± 3.98 | 33.82 ± 2.59 | 36.21 ± 5.21 | 30.38 ± 6.52 |  |
|  | 600 | 39.95 ± 3.52 | 29.91 ± 4.29 | 35.71 ± 5.22 | 28.64 ± 6.47 |  |
|  | 700 | 39.33 ± 3.57 | 30.76 ± 4.66 | 33.34 ± 5.10 | 28.05 ± 6.77 |  |
|  | 800 | 37.01 ± 3.57 | 30.38 ± 4.82 | **31.54 ± 5.14** | **36.45 ± 3.65\*** |  |
|  | 900 | 33.55 ± 3.82 | 29.58 ± 5.29 | **28.86 ± 5.33** | **33.92 ± 4.06\*** |  |
|  | 1000 | 34.14 ± 4.92 | 28.08 ± 5.47 | **26.90 ± 5.19** | **32.09 ± 4.35\*** |  |
|  | 1100 | 26.62 ± 4.68 | 25.35 ± 6.04 | 23.58 ± 5.40 | 29.68 ± 5.03 |  |
|  | 1200 | 22.43 ± 4.01 | 24.74 ± 5.59 | 22.75 ± 3.77 | 28.32 ± 4.51 |  |
| Peak Amplitude (mV) | 100 | - | - | 108.47 ± 0.00 | 84.19 ± 0.00 | Current Injected<br>P<0.0001<br>Treatment x<br>Current Injected<br>P=0.0007 |
|  | 200 | 123.91 ± 4.55 | - | 122.18 ± 10.40 | 111.43 ± 13.94 |  |
|  | 300 | 129.56 ± 3.05 | 125.72 ± 1.49 | 131.40 ± 6.02 | 118.36 ± 8.57 |  |
|  | 400 | 131.51 ± 4.39 | 121.42 ± 4.88 | 126.89 ± 4.99 | 126.28 ± 7.04 |  |
|  | 500 | 132.60 ± 4.04 | 126.74 ± 2.33 | 128.45 ± 5.24 | 122.41 ± 6.48 |  |
|  | 600 | 132.05 ± 3.56 | 122.45 ± 4.12 | 127.95 ± 5.26 | 120.55 ± 6.45 |  |
|  | 700 | 131.43 ± 3.61 | 123.17 ± 4.46 | 125.58 ± 5.14 | 119.96 ± 6.75 |  |
|  | 800 | 129.11 ± 3.64 | 122.80 ± 4.61 | 123.78 ± 5.18 | 127.79 ± 3.54 |  |
|  | 900 | 125.65 ± 3.84 | 122.00 ± 5.09 | 121.10 ± 5.37 | 125.25 ± 3.90 |  |
|  | 1000 | 122.73 ± 3.95 | 120.50 ± 5.28 | 119.14 ± 5.22 | 123.43 ± 4.18 |  |
|  | 1100 | 118.73 ± 4.74 | 117.76 ± 5.85 | 115.82 ± 5.43 | 121.01 ± 4.84 |  |
|  | 1200 | 114.54 ± 4.10 | 117.16 ± 5.41 | 114.99 ± 3.77 | 119.39 ± 4.42 |  |
| Threshold (mV) | 700 pA | -49.07 ± 1.75 | -52.32 ± 3.94 | -49.6 ± 1.68 | -51.82 ± 3.81 |  |
| Rheobase (pA) | - | 407.41 ± 21.54 | 579.63 ± 62.1 | 325.93 ± 37.6 | 433.33 ± 50 | Treatment<br>P=0.003<br>Sex P=0.03 |

Overshoot amplitude showed significant effects of current injected (Three-way RM ART ANOVA; *F*(11,257)=16.7, *p*<0.0001) and a treatment × current interaction (*F*(11,257)=2.96, *p*=0.001). Males had a significant treatment × current interaction (Two-way RM ART ANOVA; *F*(11,132)=3.9, *p*<0.0001). Planned comparisons using ART contrasts with Šidák correction revealed significant differences between saline and EtOH-treated males at high current steps: 800 pA (*t*(9.01) = −2.27, *p*=0.049), 900 pA (*t*(9.01) = −2.27, *p*=0.049), and 1000 pA (*t*(9.01) = −2.27, *p*=0.050).

Peak amplitude also exhibited significant effects of current injected (Three-way RM ART ANOVA; *F*(11,258)=21.4, *p*<0.0001) and a treatment × current interaction (*F*(11,258)=3.05, *p*=0.0007). When analyzed by sex, males showed a significant treatment × current interaction (*F*(11,132)=4.3, *p*<0.0001), whereas females did not exhibit the same effects.

The threshold of the first action potential at 700 pA, the lowest current step at which all cells fired, was also analyzed and did not differ by treatment or sex (Supplemental Table 1).

Rheobase, defined as the minimal current required to elicit an action potential, showed significant effects (Two-way ART ANOVA; Treatment: *F*(1,32)=10.4, *p*=0.003; Sex: *F*(1,32)=5.1, *p*=0.03). Planned comparisons revealed a significant difference between EtOH-exposed females and saline males (ART contrasts with Šidák correction: (*t*(32) = 3.69, *p*=0.005).

Overall, these electrophysiological findings indicate that P7 EtOH exposure reduces action potential firing in VGLUT2+ bursting neurons in females, whereas in males, EtOH exposure primarily affects action potential overshoot amplitude during high-current injections.

### 3.3 P7 EtOH exposure reduced the burstiness of VGLUT2+ neurons in females

We then analyzed how many action potentials were fired within a burst (**Figure 5A-B**). Using a three-way RM ART ANOVA, there were significant effects of treatment (*F*(1,26)=10.5, *p*=0.003), and current injected (*F*(11,334)=42.3, *p*=<0.0001). When analyzed separately by sex, females exhibited a significant treatment effect (Two-way RM ART ANOVA; (*F*(1,13)=5.509, *p*=0.0354), and a treatment x current injected interaction (*F*(11,171)=2.6, *p*=0.004). Males also exhibited a significant treatment effect (Two-Way RM ART ANOVA; *F*(1,13)=5.517, *p*=0.0353), with no significant interaction (*F*(11,163)=0.95, *p*=0.5) (**Supplementary Table 1**).

**Figure 5.**
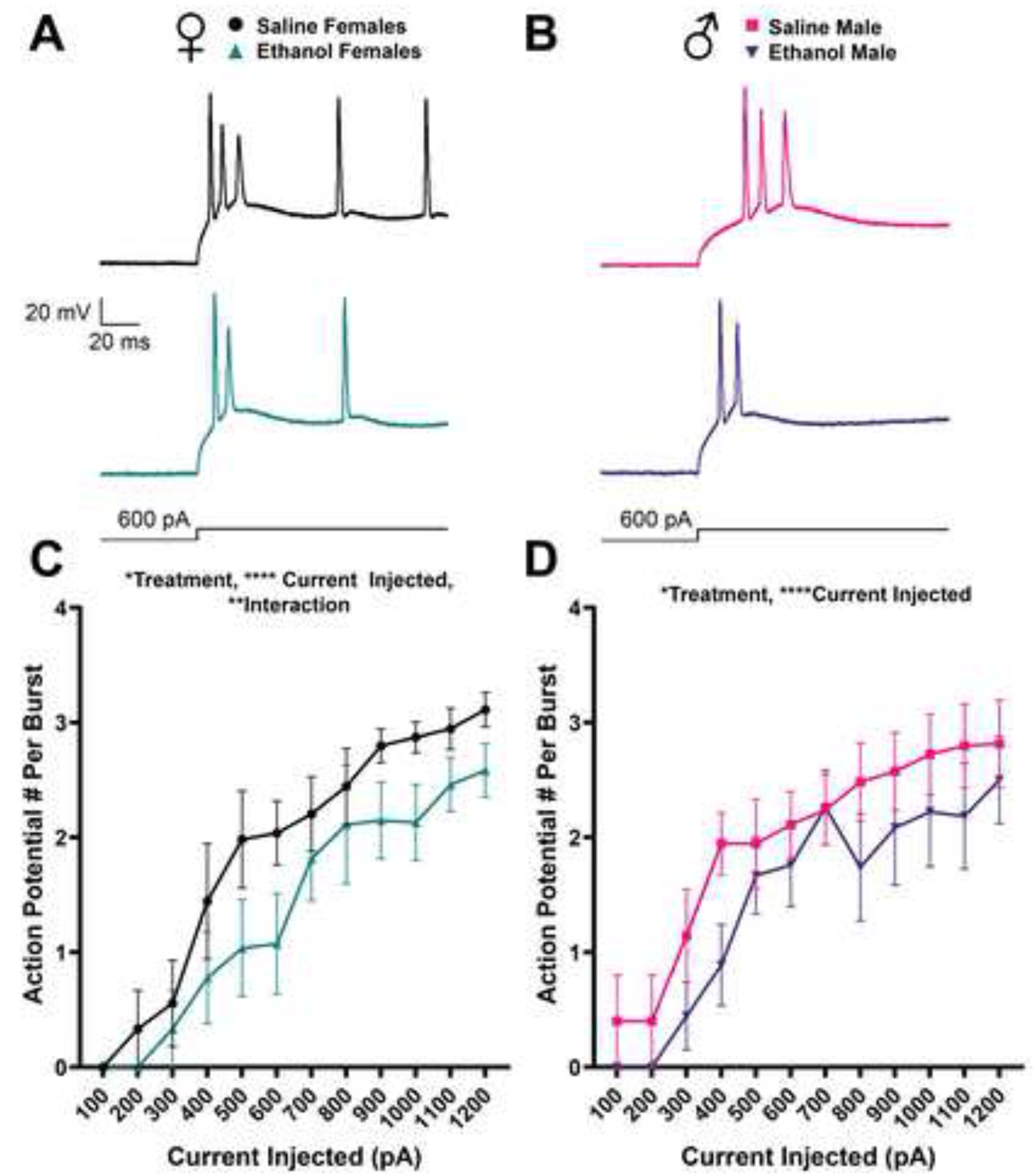
P7 ethanol exposure reduces burst strength in VGLUT2+ pyramidal neurons. A) Representative traces showing burst firing evoked by a 600-pA current injection in saline (top, black) and EtOH-treated female mice (bottom, turquoise). Scale bar = 20 ms × 20 mV. B) Representative traces from saline (top, pink) and EtOH-treated (bottom, purple) male mice. C) Summary graph illustrating the number of action potentials per burst in female mice across current injection steps. Data were analyzed using two-way ART ANOVA, revealing significant effects of treatment (\**p* = 0.0354), current injected (\*\*\*\**p* < 0.0001), and a treatment × current injected interaction (\*\**p* = 0.004). D) Equivalent analysis in males. Two-way ART ANOVA revealed significant effects of treatment (\**p* = 0.0353) and current injected (\*\*\*\**p* < 0.0001), with no significant interaction. Female control, *n* = 9 mice from 4 litters; female EtOH, *n* = 9 mice from 5 litters; male control, *n* = 9 mice from 3 litters; male EtOH, *n* = 9 mice from 4 litters.

To further characterize firing patterns in dSUB neurons, we classified recorded cells by bursting phenotype. Previous studies have identified three firing patterns in dSUB neurons: regular firing, weak bursting, and strong bursting (Rayi & Kaphzan, 2021) (**Figure 6A**). Weak bursting was defined as one burst per 500 ms current step, strong bursting as more than one burst per step, and regular firing as the absence of bursting. We compared the proportions of these firing phenotypes across sex and treatment groups. Fisher’s exact test revealed a significant treatment effect in females (*p*=0.01) and a significant sex effect in EtOH-exposed animals (*p*=0.048) (**Figure 6B**). In EtOH-exposed females, 4 of 21 recorded cells exhibited a regular firing phenotype, whereas all cells from saline-treated females displayed weak or strong bursting patterns. In males, only weak bursting neurons were observed regardless of treatment. Together, these results indicate that P7 EtOH exposure reduces number of action potentials per burst, even removing bursting in some VGLUT2+ pyramidal neurons of the dSUB in EtOH-exposed females.

**Figure 6.**
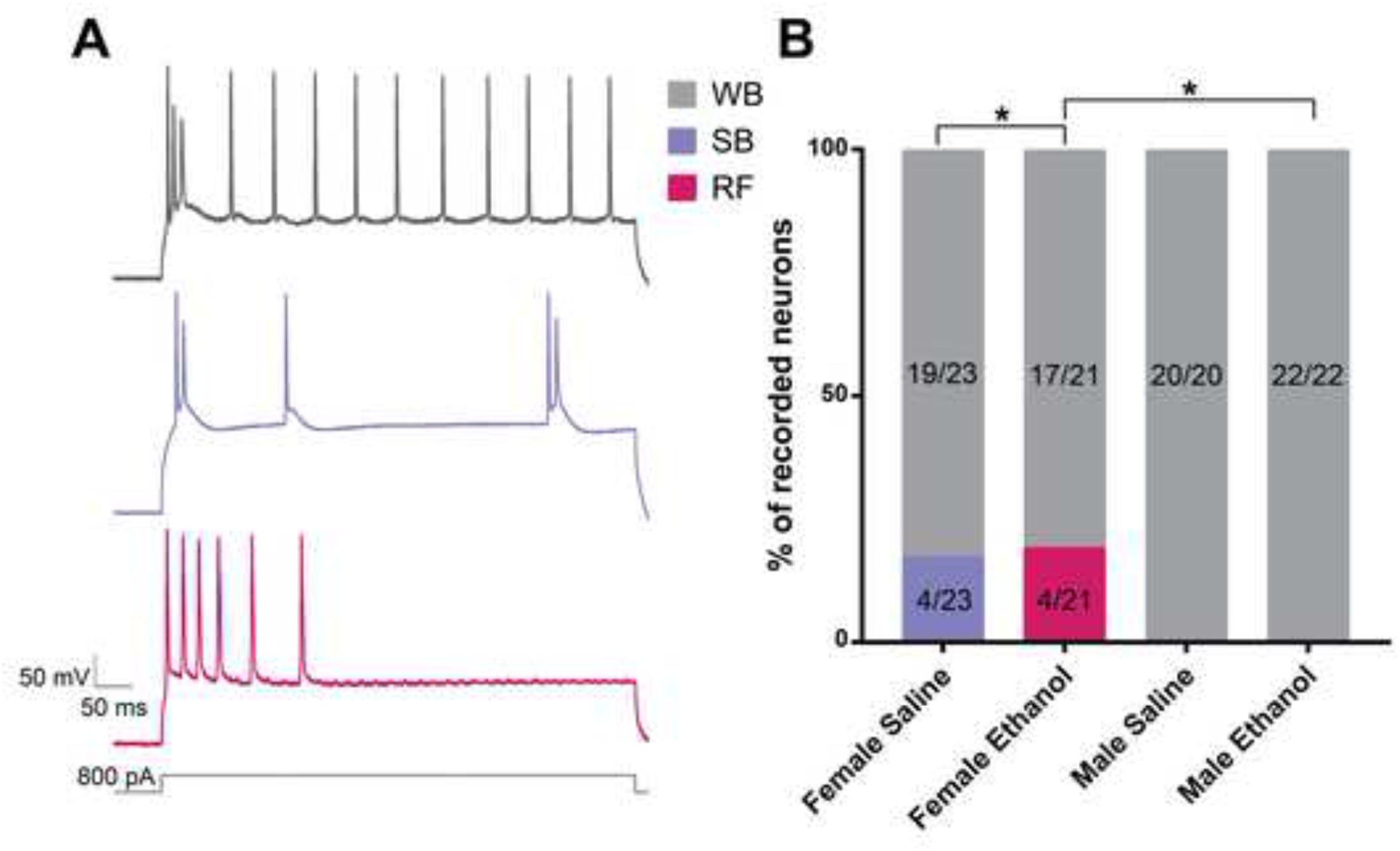
P7 ethanol exposure alters burst firing phenotypes in females. A) Representative traces illustrating three burst firing phenotypes observed in VGLUT2+ dorsal subicular neurons: weak bursting (WB; one burst per 500 ms step, grey), strong bursting (SB; two to three bursts, purple), and regular firing (RF; non-bursting, pink). Traces were evoked with an 800-pA current injection. Scale bar = 50 ms × 50 mV. B) Summary graph showing the percentage of recorded neurons classified into each burst phenotype across sex and treatment groups. Fisher’s exact test revealed a significant treatment effect in females (\**p* = 0.0125) and a significant sex difference in EtOH-treated mice (\**p* = 0.0485). Numbers within bars represent the number of neurons in each category relative to the total recorded neurons per group. Female control, *n* = 9 mice from 4 litters; female EtOH, *n* = 9 mice from 5 litters; male control, *n* = 9 mice from 3 litters; male EtOH, *n* = 9 mice from 4 litters.

## 4. Discussion

Our findings demonstrate sex-specific effects of TTAAE on the dSUB, with reduced neuronal density in males and increased VGLUT2+ cell density in females, alongside reduced neuronal excitability in both sexes and decreased bursting in females. TTAAE has been consistently shown to induce widespread apoptosis across the developing brain, which is region and cell type specific. The dSUB has emerged as a region particularly vulnerable to EtOH’s neurotoxic effects (Farber et al., 2010; Ikonomidou et al., 2000; Olney et al., 2002; Smiley et al., 2023; Wozniak et al., 2004). Consistent with our previous study, NeuN density was decreased only in males (Lopez et al., 2025); however, other studies have reported that TTAAE reduces cell densities in both sexes (Smiley et al., 2023). This discrepancy may reflect methodological differences, as we used transgenic mouse lines as well as anti-NeuN labeling specifically to quantify neurons rather than total cell populations. Prior studies suggest that TTAAE selectively affects specific neuronal subtypes rather than causing uniform cell loss. For example, a previous study reported reductions in parvalbumin+ interneurons following TTAAE (Sadrian et al., 2014). While our previous work found no change in total GABAergic interneuron density, supporting the idea that EtOH may target specific neuron subtypes in a region and cell type dependent matter.

In contrast to our initial hypothesis, VGLUT2+ cell density was increased in females following TTAAE, despite evidence of overall neuronal vulnerability within the dSUB (Farber et al., 2010; Ikonomidou et al., 2000; Olney et al., 2002; Smiley et al., 2023; Wozniak et al., 2004). Given that VGLUT2+ pyramidal neurons are critical for hippocampal output and memory-related processing, these findings suggest sex-specific reorganization of the dSUB circuitry rather than uniform degeneration. One potential explanation for the increase in VGLUT2+ cells in females is a compensatory response to reduced inhibitory tone (Lopez et al., 2025; Sadrian et al., 2014). Parvalbumin+ interneurons are reduced in the dSUB (Sadrian et al., 2014), though it remains unclear whether this loss is region specific along the proximal-distal axis. Combined with increased excitatory neurons, this may lead to hyperexcitability in the dSUB in females following TTAAE—a region critical for memory and seizure propagation (De Melo et al., 2023; Fei et al., 2022). Importantly, individuals with FASDs have an increased risk of epilepsy (Bell et al., 2010).

Our findings reveal a decrease in action potential firing in VGLUT2+ bursting neurons in adolescent females, indicating long-lasting impairments in intrinsic excitability following TTAAE. Several other labs have observed that postnatal EtOH exposure during the third-trimester equivalent has resulted in decreased action potential potentials from various neuronal populations such as dopaminergic neurons of the ventral tegmental area, layer V pyramidal neurons in the somatosensory cortex, and layer VI pyramidal neurons in the medial prefrontal cortex (Granato et al., 2012; Louth et al., 2016; Wang et al., 2006). Our laboratory has also shown that TTAAE reduced instantaneous action potential frequency in retrosplenial-projecting anterodorsal thalamic nucleus neurons (Bird et al., 2023), and action potential frequency of fast spiking GABAergic interneurons of the dSUB in females (Lopez et al., 2025). In contrast, males did not exhibit a significant reduction in overall firing, suggesting sex-dependent vulnerability of subicular neurons. Although EtOH exposure significantly reduced action potential overshoot amplitude at high current steps in males. A previous study from our laboratory has also shown that TTAAE reduces overshoot amplitude in retrosplenial-projecting anterodorsal thalamic nuclei neurons (Bird et al., 2023). Future work should include an investigation of the effect of TTAAE on Na^+^ and K^+^ channels that govern action potential amplitude. These results indicate TTAAE alters dSUB neuronal function in both sexes, but through distinct physiological mechanisms.

Another finding of this study is that TTAAE reduces burst firing in VGLUT2+ neurons; both females and males exhibited fewer action potentials per burst. Burst firing is a defining feature of subicular pyramidal neurons and is critical for information transfer from the hippocampus to downstream targets, including but not limited to the retrosplenial and entorhinal cortices (Simonnet & Brecht, 2019). The observed reduction in bursting—and the emergence of regular-spiking neurons in EtOH-exposed females—suggests a disruption in the intrinsic firing mode of these cells. Given that bursting supports the encoding of spatial and contextual information (Simonnet & Brecht, 2019), this shift toward reduced or absent bursting likely results in functional impairment in hippocampal output. Notably, all neurons recorded from control females exhibited bursting, whereas a subset of EtOH-exposed females lost this capacity entirely. This raises the possibility that VGLUT2 expression may not be an entirely reliable marker of bursting neurons. In the four EtOH-treated females, the labeled population may have included neurons exhibiting regular firing properties, rather than exclusively burst-firing cells. Despite these functional impairments, passive membrane properties remained unchanged, suggesting that TTAAE does not broadly disrupt basic cellular integrity but instead targets active conductances that underlie excitability and burst generation. Previous work identified a Ca^2+^ tail current in subicular pyramidal neurons that regulates burst firing. During an action potential, voltage-gated Ca^2+^ channels are activated and then deactivate slowly during repolarization, generating an afterdepolarization that can trigger subsequent action potentials within a burst (Jung et al., 2001). A more recent study identified T-type Ca^2+^ channels as necessary for bursting in the SUB, as pharmacological inhibition of these channels caused burst-firing neurons to become regular-spiking (Joksimovic et al., 2023). This study also found that Ca_V_3.1 channels are likely the most highly expressed isoform of T-type Ca^2+^ channels in the rat SUB. Given that both sexes exhibited a reduction in action potentials per burst, it is possible these T-type Ca^2+^ channels may have reduced efficacy or expression, potentially even more so in females.

The dSUB serves as a major output node of the hippocampal formation, and disruptions in its firing properties are likely to have downstream consequences for memory, spatial navigation, and network synchronization. The observed reduction in bursting may impair the ability of subicular neurons to effectively drive target regions, contributing to the cognitive deficits associated with FASD (Lopez et al., 2026). Ethanol exposure during the brain growth spurt specifically has been shown by numerous laboratories to impair memory, especially spatial memory (Girard et al., 2000; Goodlett & Johnson, 1997; Hamilton et al., 2010; Morningstar et al., 2024; Wozniak et al., 2004). Altered excitability in the dSUB may increase susceptibility to epileptiform activity, as preclinical models identify distinct mechanisms underlying seizure initiation and propagation in this brain region (Fei et al., 2021). Our previous work showed that inhibitory post synaptic potentials within the dSUB did not change due to EtOH exposure, even though the GABAergic interneurons had a reduction in firing (Lopez et al., 2025). The increase in VGLUT2+ cells could anticipate hyperexcitability in the dSUB, except that the VGLUT2+ cells have a reduction in action potential firing and bursting, which could be a homeostatic compensatory effect. Identifying specific receptors mediating functional deficits observed in the dSUB will be key in future therapeutics for individuals with FASDs. Additional clarity is warranted on how these intrinsic firing properties reflect on the local circuit within the dSUB as well as broader circuits and how that relates to behavioral deficits seen in the TTAAE model.

## 5. Conclusion

In summary, our findings demonstrate that P7 EtOH exposure produces long-lasting, sex-specific disruptions in the firing properties of VGLUT2+ subicular neurons, characterized by: reduced action potential firing in females, reduced action potential amplitude in males, decreased bursting, and loss of burst phenotypes in females. These results highlight the dSUB as a critical and vulnerable target of developmental alcohol exposure. Future studies should delineate the molecular mechanisms underlying altered burst firing—particularly the role of specific ion channels—and determine whether these changes arise from cell-intrinsic modifications or compensatory circuit-level adaptations. In addition, it will be important to assess the behavioral consequences of disrupted subicular output, especially with respect to memory function and seizure susceptibility.

## Acknowledgements

Author Roles: Katalina M. Lopez (conceptualization, data curation, formal analysis, investigation, visualization, writing—original draft and review & editing), Hyemin Choi (formal analysis, validation, writing - review & editing), Adeline Feng (formal analysis, validation, writing - review & editing), Lourdes Cazares (formal analysis, validation, writing - review & editing), Javier Kelly Roman (formal analysis, validation, writing - review & editing), Glenna J. Chavez (formal analysis, validation, writing - review & editing), Melissa Molina Garcia (formal analysis, validation, writing - review & editing), C. Fernando Valenzuela (conceptualization; data curation; formal analysis; funding acquisition; project administration; supervision; visualization; writing - original draft and review & editing)

In preparation of writing this manuscript, authors used ChatGPT and Grammarly to improve the readability. Authors reviewed and edited the content as needed and take full responsibility for the content of the publication.

## Disclosure/Funding

The authors have no competing interest to declare

The National Instituted of Health supported this work under grant R01 AA015614 and P50 AA022534 (CFV). KML is supported by T32 AA014127. The slide scanner at the Center for Brain Recovery and Repair is supported by grant P20GM109089. AF and HC are supported by the Combined BA/MD Program funded by the New Mexico State Legislature.

## Notes

### Competing Interest Statement

The authors have declared no competing interest.

## References

Astley, S. J., Aylward, E. H., Olson, H. C., Kerns, K., Brooks, A., Coggins, T. E., Davies, J., Dorn, S., Gendler, B., Jirikowic, T., Kraegel, P., Maravilla, K., & Richards, T. (2009). Magnetic Resonance Imaging Outcomes From a Comprehensive Magnetic Resonance Study of Children With Fetal Alcohol Spectrum Disorders. Alcoholism: Clinical and Experimental Research, 33(10), 1671–1689. 10.1111/j.1530-0277.2009.01004.x

Bankhead, P., Loughrey, M. B., Fernández, J. A., Dombrowski, Y., McArt, D. G., Dunne, P. D., McQuaid, S., Gray, R. T., Murray, L. J., Coleman, H. G., James, J. A., Salto-Tellez, M., & Hamilton, P. W. (2017). QuPath: Open source software for digital pathology image analysis. Scientific Reports, 7(1), 16878. 10.1038/s41598-017-17204-5

Baset, A., & Huang, F. (2024). Shedding light on subiculum’s role in human brain disorders. Brain Research Bulletin, 214, 110993. 10.1016/j.brainresbull.2024.110993

Bell, S. H., Stade, B., Reynolds, J. N., Rasmussen, C., Andrew, G., Hwang, P. A., & Carlen, P. L. (2010). The remarkably high prevalence of epilepsy and seizure history in fetal alcohol spectrum disorders. *Alcoholism*, Clinical and Experimental Research, 34(6), 1084–1089. 10.1111/j.1530-0277.2010.01184.x

Bird, C. W., Barber, M. J., Post, H. R., Jacquez, B., Chavez, G. J., Faturos, N. G., & Valenzuela, C. F. (2020). Neonatal ethanol exposure triggers apoptosis in the murine retrosplenial cortex: Role of inhibition of NMDA receptor-driven action potential firing. Neuropharmacology, 162, 107837. 10.1016/j.neuropharm.2019.107837

Bird, C. W., Chavez, G. J., Barber, M. J., & Valenzuela, C. F. (2021). Enhancement of parvalbumin interneuron-mediated neurotransmission in the retrosplenial cortex of adolescent mice following third trimester-equivalent ethanol exposure. Scientific Reports, 11(1), 1716. 10.1038/s41598-021-81173-z

Bird, C. W., Mayfield, S. S., Lopez, K. M., Dunn, B. R., Feng, A., Roberts, B. T., Almeida, R. N., Chavez, G. J., & Valenzuela, C. F. (2023). Binge-like ethanol exposure during the brain growth spurt disrupts the function of retrosplenial cortex-projecting anterior thalamic neurons in adolescent mice. Neuropharmacology, 241, 109738. 10.1016/j.neuropharm.2023.109738

Böhm, C., Peng, Y., Maier, N., Winterer, J., Poulet, J. F. A., Geiger, J. R. P., & Schmitz, D. (2015). Functional Diversity of Subicular Principal Cells during Hippocampal Ripples. The Journal of Neuroscience, 35(40), 13608–13618. 10.1523/JNEUROSCI.5034-14.2015

Cembrowski, M. S., Phillips, M. G., DiLisio, S. F., Shields, B. C., Winnubst, J., Chandrashekar, J., Bas, E., & Spruston, N. (2018). Dissociable Structural and Functional Hippocampal Outputs via Distinct Subiculum Cell Classes. Cell, 174(4), 1036. 10.1016/j.cell.2018.07.039

De Melo, M. B., Daldegan-Bueno, D., Favaro, V. M., & Oliveira, M. G. M. (2023). The subiculum role on learning and memory tasks using rats and mice: A scoping review. Neuroscience & Biobehavioral Reviews, 155, 105460. 10.1016/j.neubiorev.2023.105460

Farber, N. B., Creeley, C. E., & Olney, J. W. (2010). Alcohol-induced neuroapoptosis in the fetal macaque brain. Neurobiology of Disease, 40(1), 200–206. 10.1016/j.nbd.2010.05.025

Fei, F., Wang, X., Wang, Y., & Chen, Z. (2021). Dissecting the role of subiculum in epilepsy: Research update and translational potential. Progress in Neurobiology, 201, 102029. 10.1016/j.pneurobio.2021.102029

Fei, F., Wang, X., Xu, C., Shi, J., Gong, Y., Cheng, H., Lai, N., Ruan, Y., Ding, Y., Wang, S., Chen, Z., & Wang, Y. (2022). Discrete subicular circuits control generalization of hippocampal seizures. Nature Communications, 13(1), 5010. 10.1038/s41467-022-32742-x

Fox, J., & Weisberg, S. (2019). An {R} Companion to Applied Regression (Third). Sage. https://www.john-fox.ca/Companion/

Gimbel, B. A., Wozniak, J. R., Mueller, B. A., Tuominen, K. A., Ernst, A. M., Anthony, M. E., de Water, E., CIFASD, & Roediger, D. J. (2025). Regional hippocampal thinning and gyrification abnormalities and associated cognition in children with prenatal alcohol exposure. Journal of Neurodevelopmental Disorders, 17(1), 5. 10.1186/s11689-025-09595-8

Girard, T. A., Xing, H. C., Ward, G. R., & Wainwright, P. E. (2000). Early postnatal ethanol exposure has long-term effects on the performance of male rats in a delayed matching-to-place task in the Morris water maze. Alcoholism, Clinical and Experimental Research, 24(3), 300–306.

Goodlett, C. R., & Johnson, T. B. (1997). Neonatal binge ethanol exposure using intubation: Timing and dose effects on place learning. Neurotoxicology and Teratology, 19(6), 435–446. 10.1016/s0892-0362(97)00062-7

Granato, A., Palmer, L. M., De Giorgio, A., Tavian, D., & Larkum, M. E. (2012). Early exposure to alcohol leads to permanent impairment of dendritic excitability in neocortical pyramidal neurons. The Journal of Neuroscience: The Official Journal of the Society for Neuroscience, 32(4), 1377–1382. 10.1523/JNEUROSCI.5520-11.2012

Hamilton, G. F., Whitcher, L. T., & Klintsova, A. Y. (2010). Postnatal binge-like alcohol exposure decreases dendritic complexity while increasing the density of mature spines in mPFC Layer II/III pyramidal neurons. Synapse, 64(2), 127–135. 10.1002/syn.20711

Hoyme, H. E., Kalberg, W. O., Elliott, A. J., Blankenship, J., Buckley, D., Marais, A.-S., Manning, M. A., Robinson, L. K., Adam, M. P., Abdul-Rahman, O., Jewett, T., Coles, C. D., Chambers, C., Jones, K. L., Adnams, C. M., Shah, P. E., Riley, E. P., Charness, M. E., Warren, K. R., & May, P. A. (2016). Updated Clinical Guidelines for Diagnosing Fetal Alcohol Spectrum Disorders. Pediatrics, 138(2), e20154256. 10.1542/peds.2015-4256

Ikonomidou, C., Bittigau, P., Ishimaru, M. J., Wozniak, D. F., Koch, C., Genz, K., Price, M. T., Stefovska, V., Hörster, F., Tenkova, T., Dikranian, K., & Olney, J. W. (2000). Ethanol-Induced Apoptotic Neurodegeneration and Fetal Alcohol Syndrome. Science, 287(5455), 1056–1060. 10.1126/science.287.5455.1056

Joksimovic, S. M., Ghodsi, S. M., Heinsbroek, J. A., Orfila, J. E., Busquet, N., Tesic, V., Valdez, R., Fine-Raquet, B., Jevtovic-Todorovic, V., Raol, Y. H., Herson, P. S., & Todorovic, S. M. (2023). CaV3.1 T-type calcium channels are important for spatial memory processing in the dorsal subiculum. Neuropharmacology, 226, 109400. 10.1016/j.neuropharm.2022.109400

Jung, H. Y., Staff, N. P., & Spruston, N. (2001). Action potential bursting in subicular pyramidal neurons is driven by a calcium tail current. The Journal of Neuroscience: The Official Journal of the Society for Neuroscience, 21(10), 3312–3321. 10.1523/JNEUROSCI.21-10-03312.2001

Kay, M., Elkin, L. A., Higgins, J. J., & Wobbrock, J. O. (2025). {ARTool}: Aligned Rank Transform for Nonparametric Factorial ANOVAs. 10.5281/zenodo.594511

Lopez, K. M., Mancero-Montalvo, R., Iturralde-Carrillo, A., Feng, A., Shaver, T., Choi, H., Jaiswal, N., Kelly-Roman, J., Bhakta, P. A., Chavez, G. J., & Valenzuela, C. F. (2025). Impact of acute binge-like ethanol exposure during the third-trimester equivalent on subicular interneurons in mice. The American Journal of Drug and Alcohol Abuse, 1–13. 10.1080/00952990.2025.2529503

Lopez, K. M., Shaver, T. L., & Fernando Valenzuela, C. (2026). Prenatal alcohol damages the subiculum, a key region for learning and memory: A narrative review. Alcohol, 133, 71–86. 10.1016/j.alcohol.2026.03.008

Louth, E. L., Bignell, W., Taylor, C. L., & Bailey, C. D. C. (2016). Developmental Ethanol Exposure Leads to Long-Term Deficits in Attention and Its Underlying Prefrontal Circuitry. eNeuro, 3(5), ENEURO.0267-16.2016. 10.1523/ENEURO.0267-16.2016

May, P. A., Chambers, C. D., Kalberg, W. O., Zellner, J., Feldman, H., Buckley, D., Kopald, D., Hasken, J. M., Xu, R., Honerkamp-Smith, G., Taras, H., Manning, M. A., Robinson, L. K., Adam, M. P., Abdul-Rahman, O., Vaux, K., Jewett, T., Elliott, A. J., Kable, J. A., … Hoyme, H. E. (2018). Prevalence of Fetal Alcohol Spectrum Disorders in 4 US Communities. JAMA, 319(5), 474–482. 10.1001/jama.2017.21896

Morningstar, M. D., Lopez, K. M., Mayfield, S. S., Almeida-Mancero, R. N., Marquez, J., Flores, A. M., Hafer, B. R., Estrada, E., Holtzman, G. A., Goranson, E. V., Reid, N. M., Aldrich, A. R., Ghatalia, D. V., Patel, J. R., Padilla, C. M., Chavez, G. J., Kelly-Roman, J., Bhakta, P. A., Valenzuela, C. F., & Linsenbardt, D. N. (2024). Connectivity of the neuronal network for contextual fear memory is disrupted in a mouse model of third-trimester binge-like ethanol exposure. Alcohol, Clinical & Experimental Research. 10.1111/acer.15503

Nakhid, D., Patel, D., McMorris, C. A., Gibbard, W. B., Tortorelli, C., Pei, J., & Lebel, C. (2023). Limbic brain subregions associated with mental health symptoms in youth with and without prenatal alcohol exposure. Alcohol: Clinical and Experimental Research, 47(11), 2033–2044. 10.1111/acer.15181

Nitzan, N., McKenzie, S., Beed, P., English, D. F., Oldani, S., Tukker, J. J., Buzsáki, G., & Schmitz, D. (2020). Propagation of hippocampal ripples to the neocortex by way of a subiculum-retrosplenial pathway. Nature Communications, 11(1), 1947. 10.1038/s41467-020-15787-8

Olney, J. W., Tenkova, T., Dikranian, K., Qin, Y.-Q., Labruyere, J., & Ikonomidou, C. (2002). Ethanol-induced apoptotic neurodegeneration in the developing C57BL/6 mouse brain. Developmental Brain Research, 133(2), 115–126. 10.1016/S0165-3806(02)00279-1

Paxinos, G., & Franklin, K. B. J. (2019). Paxinos and Franklin’s The mouse brain in stereotaxic coordinates (Fifth edition). Academic Press, an imprint of Elsevier.

Popova, S., Charness, M. E., Burd, L., Crawford, A., Hoyme, H. E., Mukherjee, R. A. S., Riley, E. P., & Elliott, E. J. (2023). Fetal alcohol spectrum disorders. Nature Reviews. Disease Primers, 9(1), 11. 10.1038/s41572-023-00420-x

R Core Team. (2024). R: A Language and Environment for Statistical Computing. https://www.R-project.org

Rayi, P. R., & Kaphzan, H. (2021). Electrophysiological Characterization of Regular and Burst Firing Pyramidal Neurons of the Dorsal Subiculum in an Angelman Syndrome Mouse Model. Frontiers in Cellular Neuroscience, 15, 670998. 10.3389/fncel.2021.670998

Roediger, D. J., Krueger, A. M., De Water, E., Mueller, B. A., Boys, C. A., Hendrickson, T. J., Schumacher, M. J., Mattson, S. N., Jones, K. L., Lim, K. O., & Wozniak, J. R. (2021). Hippocampal subfield abnormalities and memory functioning in children with fetal alcohol Spectrum disorders. Neurotoxicology and Teratology, 83, 106944. 10.1016/j.ntt.2020.106944

Sadrian, B., Lopez-Guzman, M., Wilson, D. A., & Saito, M. (2014). Distinct neurobehavioral dysfunction based on the timing of developmental binge-like alcohol exposure. Neuroscience, 280, 204–219. 10.1016/j.neuroscience.2014.09.008

Simonnet, J., & Brecht, M. (2019). Burst Firing and Spatial Coding in Subicular Principal Cells. The Journal of Neuroscience: The Official Journal of the Society for Neuroscience, 39(19), 3651–3662. 10.1523/JNEUROSCI.1656-18.2019

Smiley, J. F., Bleiwas, C., Marino, B. M., Vaddi, P., Canals-Baker, S., Wilson, D. A., & Saito, M. (2023). Estimates of total neuron number show that neonatal ethanol causes immediate and lasting neuron loss in cortical and subcortical areas. Frontiers in Neuroscience, 17, 1186529. 10.3389/fnins.2023.1186529

Solar, K. G., Treit, S., & Beaulieu, C. (2022). High-resolution diffusion tensor imaging identifies hippocampal volume loss without diffusion changes in individuals with prenatal alcohol exposure. Alcoholism: Clinical and Experimental Research, 46(7), 1204–1219. 10.1111/acer.14857

Wang, J., Haj-Dahmane, S., & Shen, R.-Y. (2006). Effects of prenatal ethanol exposure on the excitability of ventral tegmental area dopamine neurons in vitro. The Journal of Pharmacology and Experimental Therapeutics, 319(2), 857–863. 10.1124/jpet.106.109041

Wickham, H., & Bryan, J. (2025). readxl: Read Excel Files. https://CRAN.R-project.org/package=readxl

Willoughby, K. A., Sheard, E. D., Nash, K., & Rovet, J. (2008). Effects of prenatal alcohol exposure on hippocampal volume, verbal learning, and verbal and spatial recall in late childhood. Journal of the International Neuropsychological Society, 14(6), 1022–1033. 10.1017/S1355617708081368

Wozniak, D., Hartman, R., Boyle, M., Vogt, S., Brooks, A., Tenkova, T., Young, C., Olney, J., & Muglia, L. (2004). Apoptotic neurodegeneration induced by ethanol in neonatal mice is associated with profound learning/memory deficits in juveniles followed by progressive functional recovery in adults. Neurobiology of Disease, 17(3), 403–414. 10.1016/j.nbd.2004.08.006

Wozny, C., Beed, P., Nitzan, N., Pössnecker, Y., Rost, B. R., & Schmitz, D. (2018). VGLUT2 Functions as a Differential Marker for Hippocampal Output Neurons. Frontiers in Cellular Neuroscience, 12, 337. 10.3389/fncel.2018.00337

